# Investigating cross-sectional and longitudinal relationships between brain structure and distinct dimensions of externalizing psychopathology in the ABCD Sample

**DOI:** 10.1101/2024.03.01.583021

**Authors:** Lee Propp, Hajer Nakua, Anne-Claude V. Bedard, Marcos Sanches, Stephanie H. Ameis, Brendan F. Andrade

## Abstract

Externalizing psychopathology in childhood is a predictor of poor outcomes across the lifespan. Children exhibiting elevated externalizing psychopathology also commonly show emotion dysregulation and callous-unemotional (CU) traits. Examining cross-sectional and longitudinal neural correlates across dimensions linked to externalizing psychopathology during childhood may clarify shared or distinct neurobiological vulnerability for psychopathological impairment later in life. We used tabulated brain structure and behavioural data from baseline, year 1, and year 2 timepoints of the Adolescent Brain Cognitive Development Study (ABCD; baseline n=10,534). We fit separate linear mixed effect models to examine whether baseline brain structures in frontolimbic and striatal regions (cortical thickness or subcortical volume) were associated with externalizing symptoms, emotion dysregulation, and/or CU traits at baseline and over a two-year period. At baseline, cortical thickness in the right rostral middle frontal gyrus and bilateral pars orbitalis was positively associated with CU traits (*β*=|0.027-0.033|, *p*_*corrected*_=0.009-0.03). Subcortical volume in the left caudate, right amygdala, and bilateral nucleus accumbens was negatively associated with emotion dysregulation (*β*=|0.026 – 0.037|, *p*_*corrected*_=<0.001-0.02). Over the two-year follow-up period, higher baseline cortical thickness in the left pars triangularis and rostral middle frontal gyrus predicted greater decreases in externalizing symptoms (F=6.33-6.94, *p*_*corrected*_=0.014). The results of the current study suggest that unique regions within frontolimbic and striatal networks may be more strongly associated with different dimensions of externalizing psychopathology. The longitudinal findings indicate that brain structure in early childhood may provide insight into structural features that influence behaviour over time.

## Introduction

Clinically significant externalizing behaviours in childhood predict development of a variety of mental health disorders in adolescence [1]. Externalizing behaviours include rule-breaking and aggressive behaviours which are among the most common reasons for referral to child mental health services [2]. Diagnostic categories such as Attention-Deficit Hyperactivity Disorder (ADHD), Oppositional Defiant Disorder (ODD), and Conduct Disorder (CD) capture a breadth of significantly impairing externalizing symptoms and associated disruptions in adaptive functioning and daily life [3]. Yet, the substantial overlap in symptom presentation between these diagnostic categories, in addition to limited knowledge on developmental trajectories of symptoms within a given category, constrains the clinical utility of diagnoses groups [4,5]. It is now increasingly recognized that similar alterations of brain structures are implicated across multiple disorders, leading to nonspecific findings [6]. As a result, recent work has emphasized the utility of investigating transdiagnostic dimensions of psychopathology such that the spectrum of symptom variation is explored in place of comparing diagnostic categories [7,8]. This approach may enable delineation of neurobiological associations across the full range of symptom expression (i.e., from the absence of symptoms to clinically relevant symptoms) to provide insight into whether or not similar brain regions are continuously associated with related symptom dimensions [9]. Given the poor mental health outcomes that childhood externalizing behaviours confer [1], a longitudinal approach is ideal to examine whether particular brain features in earlier childhood are associated with progression of symptoms.

Efforts to identify subgroups of children with externalizing behaviours using dimensions of psychopathology have reported that those with high levels of callous-unemotional (CU) traits (i.e., low levels of guilt, empathy, and care for others) are at the greatest risk for later developing severe conduct problems [10,11]. It remains unclear whether emotion dysregulation (e.g., emotional lability, irritability, and surgency) is present in some children with high externalizing behaviours and high CU traits [12]. Some studies argue that a lack of emotional arousal, or an overregulation of emotions, is thought to underlie the ‘covert’ aggression and behaviours (e.g., lying) observed in those with high CU traits [13,14]. However, a recent systematic review found evidence of significant emotion dysregulation in youth with externalizing behaviours and high CU traits [15].

Despite the emphasis on dimensional frameworks of psychopathology [16], there are limited studies that consider the neurobiological correlates of externalizing behaviours, emotion dysregulation, and CU traits in the same study sample [15], leading to inconsistent results [17,18]. However, several studies have implicated common cortical and subcortical regions across these externalizing dimensions. Included are many cortico-limbic structures that are key in mediating the adaptive regulation of emotion and related processes, such as the anterior cingulate cortex (ACC) [19,20], insula [21,22], amygdala [23,24], dorsolateral prefrontal cortex (dlPFC) [25], and ventrolateral prefrontal cortex (vlPFC) [26,27]. Investigating the neural correlates of externalizing symptoms, CU traits, and emotion dysregulation during mid-to-late childhood could provide insight into the risk of emerging psychopathology in adolescence and early adulthood given that this period is critical for brain maturation and structural reorganization [28,29].

The present study had two aims. First, to investigate cross-sectional relationships between cortical thickness/subcortical volume and externalizing symptoms, CU traits, and emotion dysregulation. Second, to investigate longitudinal trajectories of whether cortical thickness and/or subcortical volumes at baseline predict externalizing symptoms, CU traits, emotion dysregulation over a 2-year period and, relatedly, if common brain regions predict these behavioural constructs over time. To address these aims, we used the Adolescent Brain Cognitive Development Study (ABCD), a large population-based sample of children and early adolescents [30,31]. Given the convergence between externalizing symptoms, emotion dysregulation, and CU traits at the behavioural level, it was predicted that similar brain regions would be associated with these three dimensions, potentially signalling common underlying brain-based vulnerability for externalizing psychopathology in early adolescence.

## Methods

### Participants

The ABCD study includes a population-based cohort of 11,878 children followed through adolescence who were recruited from 21 sites across the United States [30]. Data collection began when the participants were 9-10 years old and continued annually or biannually. Children were recruited via presentations and emails to parents of children in schools around each site. Participants were excluded from the ABCD study if they had MRI contraindications, no English fluency, uncorrected vision, hearing impairments, major neurological disorders, an extremely preterm birth, low birth weight, birth complications, or unwillingness to complete assessments. The current study used data from the fourth annual release (DOI:10.15154/1523041). From the complete participant sample, we excluded those with poor brain imaging quality, missing T1-weighted scans, and/or missing behavioural measures (see Figure S1 for consort diagram and details of exclusion). Following these exclusions, the current study included 10,534 participants from the ABCD baseline sample, 9,962 from the 1-year follow-up, and 9,253 from the 2-year follow up. The demographic characteristics of the participants across timepoints are summarized in Table 1.

**Table 1.** Participant Demographics. Abbreviations: HS, high school; GED, general education development certificate

| Characteristic | Timepoint |  |  |
| --- | --- | --- | --- |
|  | Baseline (n=10534) | 1y follow up (n=9962) | 2y follow up (n=9253) |
|  | Mean (SD) | Mean (SD) | Mean (SD) |
| Age (years) | 9.92 (0.63) | 10.93 (0.65) | 12.01 (0.66) |
|  | N (%) | N (%) | N (%) |
| Sex at birth (male) | 5498 (52.2) | 5212 (52.3) | 4849 (52.4) |
|  | Mean (SD) | Mean (SD) | Mean (SD) |
| CBCL Subscales (t-scores) |  |  |  |
| Aggressive behaviours | 52.82 (5.51) | 52.54 (5.22) | 52.35 (4.42) |
| Attention problems | 53.85 (6.14) | 53.63 (5.90) | 53.43 (5.58) |
| Anxious/depressed | 53.5 (5.99) | 53.52 (5.98) | 53.20 (5.81) |
| ABCD Callous-Unemotional score | 0.90 (1.39) | 0.97 (1.41) | 0.97 (1.44) |
| K-SADS Externalizing symptoms composite | 0.85 (2.3) | 0.83 (2.28) | 0.72 (2.11) |
|  | N (%) | N (%) | N (%) |
| Race/ethnicity |  |  |  |
| White | 5533 (52.5) | 5358 (53.8) | 3338 (36.1) |
| Black | 1541 (14.6) | 1370 (13.8) | 657 (7.1) |
| Asian | 220 (2.1) | 211 (2.1) | 121 (1.3) |
| Hispanic | 2140 (20.3) | 1972 (19.8) | 1117 (12.1) |
| Other | 1099 (10.4) | 1031 (10.3) | 573 (6.2) |
| No response | 1 (>0.01) | 20 (0.2) | 3447 (37.3) |
| Psychiatric Medications |  |  |  |
| Stimulants | 776 (7.4) | 740 (7.4) | 690 (7.5) |
| α-2-Agonists | 206 (2.0) | 202 (2.0) | 204 (2.2) |
| Atomoxetine | 40 (0.4) | 35 (0.4) | 33 (0.4) |
| Antipsychotics | 62 (0.6) | 63 (0.6) | 69 (0.7) |
| Antidepressants | 196 (1.9) | 242 (2.4) | 285 (3.1) |
| Parent Education |  |  |  |
| < HS Diploma | 504 (4.8) |  |  |
| HS Diploma/GED | 979 (9.3) |  |  |
| Some College | 2732 (25.9) |  |  |
| Bachelor | 2683 (25.5) |  |  |
| Post Graduate Degree | 3624 (34.4) |  |  |
| Household Income |  |  |  |
| <50k | 2827 (26.8) |  |  |
| ≥ 50k & <100k | 2744 (26.0) |  |  |
| ≥ 100k | 4082 (38.8) |  |  |
| No response | 881 (8.4) |  |  |

### Measures

#### Structural MRI

The T1-weighted imaging protocol has been detailed in prior ABCD publications [31,32]. T1-weighted scans were collected and processed by the Data Analysis Informatics Resource Center (DAIRC) based on standardized ABCD protocols (for details see [32]). Cortical thickness and subcortical volume segmentation was performed using FreeSurfer v5.3.0 [33,34]. All T1-weighted scans were examined by trained visual raters who recommended that scans be excluded if they contained artefacts or were unable to be processed correctly. Participants included in the current study passed the DAIRC visual quality control (QC) criteria.

Cortical thickness of eight regions and subcortical volume from four regions in the Desikan-Killiany Atlas parcellations [35] were used as morphology measures in the current study. These regions were selected as they have been strongly linked to externalizing psychopathology, including CU traits and emotion dysregulation in prior work (see Table S1).

#### Externalizing Symptoms

Externalizing symptoms were indexed by the Kiddie-Structured Assessment for Affective Disorders and Schizophrenia (K-SADS) externalizing symptoms score. The K-SADS for Diagnostic and Statistical Manual of Mental Disorders, Fifth Edition (DSM-5) [36] is a semi-structured interview used to measure psychopathology with strong psychometric properties (*α*= 0.88) [37,38]. The computerized version was self-administered by parents of study participants. As has been done in previous ABCD studies with other symptom domains [39,40], we measured externalizing symptoms using a composite derived by summation of present ODD and CD symptoms (i.e., total number of symptoms endorsed; possible scores range from 0-25). We did not use the externalizing subscale from the Child Behaviour Checklist-Parent Report (CBCL) due to overlap of the aggressive behaviour syndrome scale included in the CBCL dysregulation subscale (see below), which was used as a measure of emotion dysregulation.

#### Emotion Dysregulation

The CBCL is a 113-item questionnaire used to comprehensively assess behavioural and emotional problems in children and adolescents [41]. It provides norm-referenced scores for internalizing and externalizing problems that are derived from eight syndrome scales. In the present study the dysregulation profile raw score (sum of anxious/depressed, attention problems, and aggressive behaviour syndrome scales) was used to measure emotion dysregulation [42].

#### Callous-Unemotional (CU) Traits

CU traits were measured using a four-item measure created and validated in previous ABCD studies [43,44] that included one item from the CBCL–Parent Report (“lack of guilt after misbehaving”) and three reverse-scored items from the prosocial behaviours subscale of the Strengths and Difficulties Questionnaire–Parent Report (SDQ; “is considerate of others feelings’”, “is helpful if someone is hurt or upset”, “offers to help others”) [45]. Reliability of the CU trait measure was acceptable (*α*= 0.75). Higher scores on this measure are indicative of elevated CU traits (i.e., more impairment).

### Statistical Analysis

There was significant skew and zero inflation of the three psychopathology dimension scores (see Table 1 for means and standard deviations of each measure). We decided to retain participants with zero-scores across these measures (i.e., no endorsement of psychopathology) to i) maximize sample size, and ii) examine the neuroanatomical correlates across the range of psychopathology presentation in the population (no psychopathology to clinically relevant symptoms). Given the large sample size of the current study (i.e., n > 500), the assumption of normally distributed outcomes is unlikely to be a major concern [46].

#### Aim 1: Cross-sectional brain-behaviour relationships

Separate linear mixed effect models were fit to examine the presence of a cross-sectional association between externalizing symptoms, emotion dysregulation, or CU traits and cortical thickness/subcortical volume regions. Fixed effect covariates included sex, age, medication status, and household income (a proxy of socioeconomic status; [47,48]). Random effects included family ID (siblings enrolled in a study were given the same family ID) and site. In each model, the psychopathology dimension of interest was the dependent variable, and the cortical or subcortical ROI was the independent variable; both variables were scaled (mean centred and standard deviation of 1) in the linear model. Each model examined the relationship between one psychopathology measure (e.g., externalizing symptoms) and one brain metric (e.g., left amygdala volume; see example below) necessitating multiple correction (at the FDR threshold) for all the models run between the three dimensions and the 24 brain metrics (48). We corrected for multiple comparisons separately for the two sets of models (i.e., cortical thickness and subcortical volume). For example, there were a total of eight models run to examine the relationship between externalizing symptoms and subcortical structures, thus, we corrected for eight p-values for this group of analyses. We continued to use this correction approach for all other groups of analyses (e.g., 16 corrected tests when running the model between externalizing symptoms and cortical structures, etc.). These analyses were run in R (version 4.1) using the *lmer* function in the *lme4* package [49,50]. See below for an example of the linear mixed effect model run to address Aim 1 (where *ijk* refers to subject *k* from family *i* at site *j*).

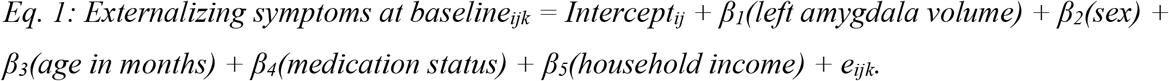

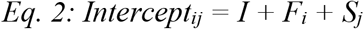

*where e*_*ijk*_*∼N(0,σ); F*_*i*_ *∼ N(0,σ*_*f*_*); S*_*j*_ *∼ N(0,σ*_*s*_*) are random effect terms related to the family (F) and site (S) respectively, and assumed to be normally distributed and uncorrelated with the fixed effect coefficients β*.

#### Aim 2: Cortical thickness and subcortical volumes as predictors of symptom trajectory

Separate linear mixed effect models were fit to examine if baseline cortical thickness and subcortical volume predicted trajectories of externalizing symptoms, emotion dysregulation, or CU traits over time. These models were also used to test the main effect of time. As in the cross-sectional models, fixed effect covariates included sex, age, medication status, and household income. Random effects included participant ID, family ID, and site. In each model, the psychopathology dimension of interest was the dependent variable, and the brain metric was the independent variable which interacted with time (baseline, year 1, year 2); both variables were scaled (mean centered with a standard deviation of 1 in the linear model). Corrections for multiple comparisons (at the FDR threshold) were carried out in a similar manner as the cross-sectional models. If the overall interaction was significant (p≤0.05) following multiple comparison correction, we interpreted it by graphing the predicted time effect (trajectory of emotion dysregulation/CU traits/externalizing symptoms) for a low, medium, and high value of the cortical thickness, defined as the first quartile, median and third quartiles. These analyses were run in R (version 4.1) using the *lmer* function in the *lme4* package. See below for an example of the linear mixed effect model run to address Aim 2 (where *ijkl* refers to subject *k* from family *i* at site *j* and time *l*).

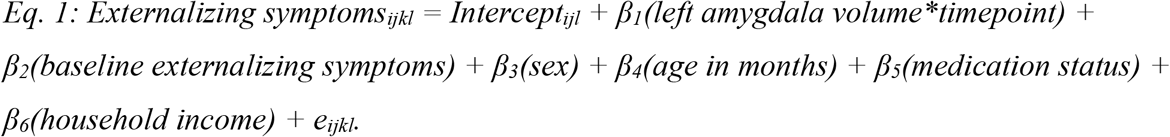

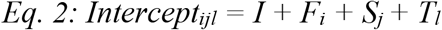

*where e*_*ijkl*_*∼N(0,σ); F*_*i*_ *∼ N(0,σ*_*f*_*); S*_*j*_ *∼ N(0,σ*_*s*_*); T*_*l*_ *∼ N(0,σ*_*s*_*) are random effect terms related to the family (F), site (S), and time (T) respectively, and assumed to be normally distributed and uncorrelated with the fixed effect coefficients β*.

### Data and Code Availability

Data for the ABCD Study are available through the National Institutes of Health Data Archive (NDA; nih.nda.gov). The participant IDs included in these analyses and details on the measures used can be found in this project’s NDA study (DOI: 10.15154/q36k-ga33). The code for the analysis can be found on GitHub (https://github.com/hajernakua/ext_psychopathology_ABCD).

## Results

*Table 1* includes the demographic and clinical characteristics of the baseline, year-1, and year-2 sample data. Overall, emotion dysregulation scores remained similar across the three timepoints, CU traits increased slightly in year-1 and year-2 compared to baseline, and externalizing symptoms decreased over time. The percentage of participants taking various medications remained relatively stable over time.

### Aim 1: Cross-sectional brain-behaviour relationships

After correcting for multiple comparisons, higher CU trait scores (greater impairment) were significantly associated with increased cortical thickness in the right rostral middle frontal gyrus (*β*=0.027, *95%CI*=0.008–0.046, *t*=2.77, *p*_*corrected*_=0.03) and the left and right pars orbitalis (left: *β*=0.033, *95%CI*=0.014–0.052, *t*=3.47, *p*_*corrected*_=0.009; right: *β*=0.027, *95%CI*=0.008– 0.046, *t*=2.82, *p*_*corrected*_=0.03). Greater emotion dysregulation was significantly associated with reduced subcortical volume in the left caudate (*β*= −0.026, *95%CI* = −0.044 –-0.007, *t*= −2.78, *p*_*corrected*_=0.02), right amygdala (*β*= −0.027, *95%CI*= −0.045 – −0.007, *t*= −2.72, *p*_*corrected*_=0.02), left and right nucleus accumbens (left: *β*= −0.024, *95%CI*= −0.04 – −0.006, *t*= −2.57, *p*_*corrected*_=0.02; right: *β*= −0.037, *95%CI*= −0.055 – −0.018, *t*= −3.92, *p*_*corrected*_ <0.001). There were no significant relationships between externalizing symptoms and cortical thickness or subcortical volume. See *Figure 1* depicting the beta weight estimates of the linear mixed models (i.e., associations between externalizing symptoms/emotion dysregulation/CU traits and cortical thickness or subcortical volume).

**Figure 1.**
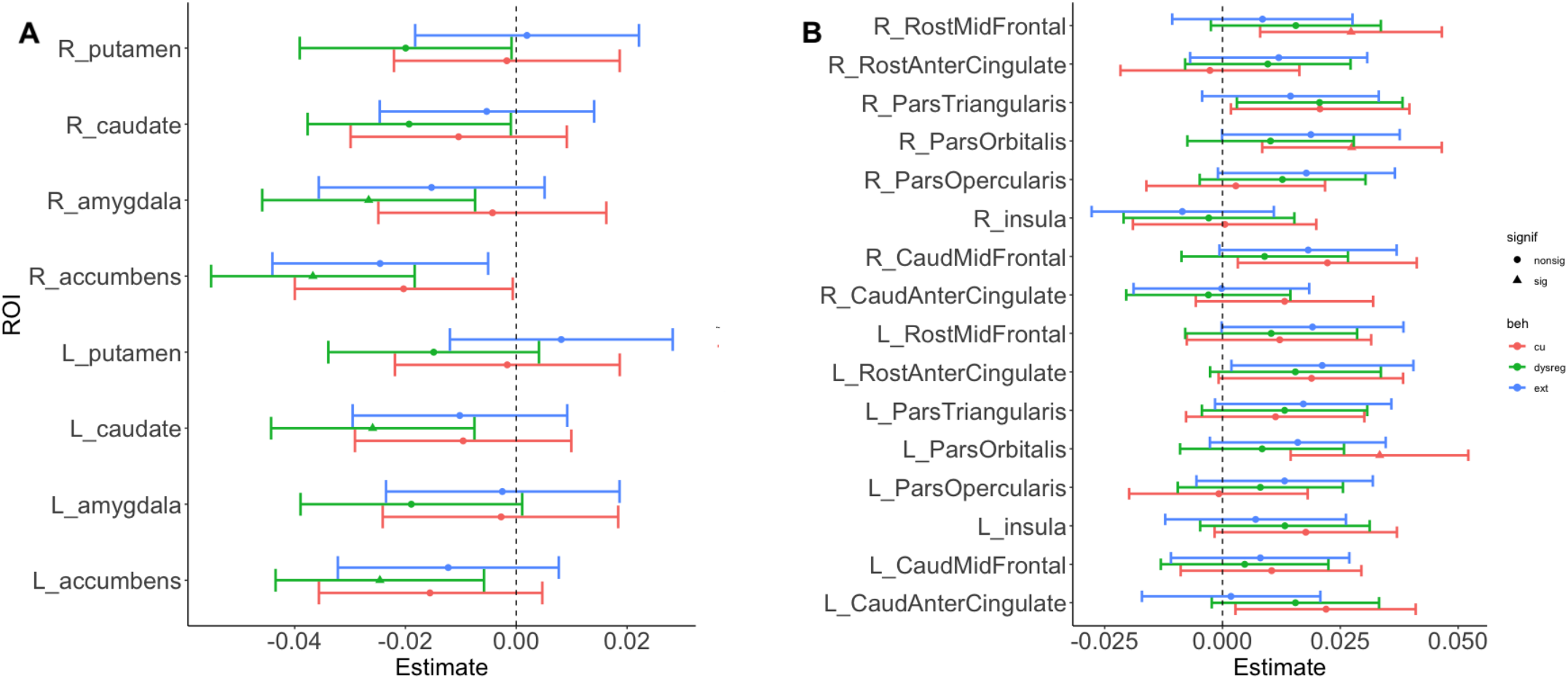
Confidence intervals of the beta weight estimates derived from the linear mixed models showing associations between externalizing symptoms/emotion dysregulation/CU traits and cortical thickness and subcortical volume. Panel A shows the models using subcortical volume ROIs as the independent variable. Panel B shows the models using cortical thickness ROIs as the independent variable. The red confidence interval line represents standardized CU trait scores, the green represents standardized emotion dysregulation scores, and the blue represents standardized externalizing symptom scores. Circles on the confidence interval bars indicate that the model did not surpass significance after correcting for multiple comparisons, and triangles indicate that those models did surpass significance after correcting for multiple comparisons. Full ROI labels are included in Table S2. The beta weight estimates across all models are low (<0.1) indicating that the effect of the brain-behaviour relationships examined are small in the current sample, as is expected given the large sample size. This data is from the baseline sample of the ABCD (ages=9-10).

### Aim 2: Cortical thickness and subcortical volumes as predictors of symptom trajectories

After correcting for multiple comparisons, lower baseline cortical thickness in the left pars triangularis (F_(2, 20576)_=6.94, *p*_*corrected*_=0.014) and left rostral middle frontal gyrus (F_(2, 20619)_=6.33, *p*_*corrected*_=0.014) moderated the trajectory of externalizing symptoms over time. *Figure 2* depicts the estimated marginal means of externalizing symptoms as a function of time (baseline, year 1, and year 2; x-axis). We found a main effect of time such that across all quartiles of cortical thickness, there was a decrease in externalizing symptoms as a function of time. That is, regardless of the variation of thickness of these regions at baseline, participants show reduction of externalizing symptoms. The significant interaction provides evidence that the decline of externalizing symptoms is greater (lower symptom expression across time) when cortical thickness of these regions is higher at baseline. There were no significant relationships between baseline cortical thickness or baseline subcortical volumes and trajectories of emotion dysregulation or CU traits over the same time period.

**Figure 2.**
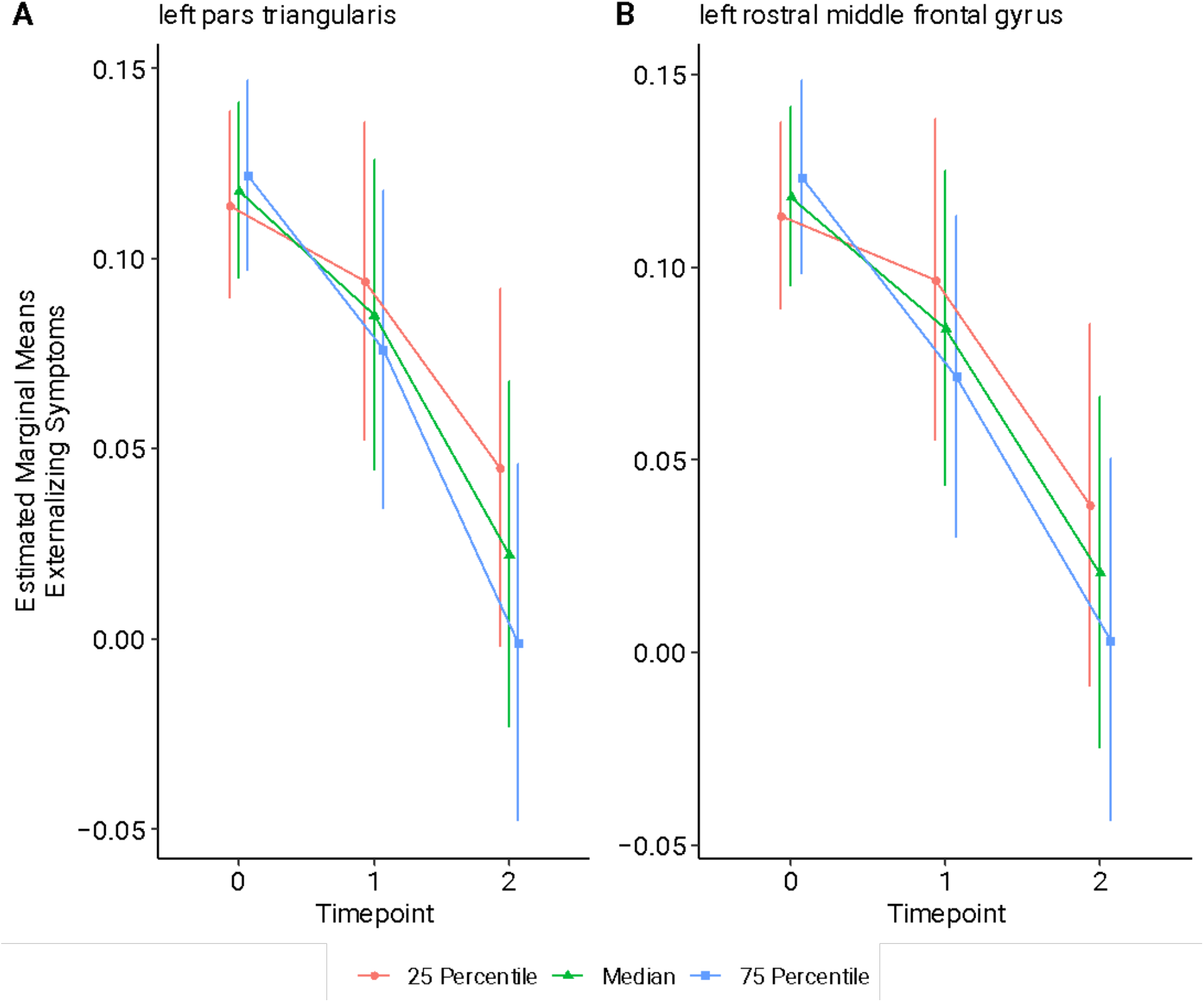
Trajectories of externalizing symptoms as predicted by baseline cortical thickness. Low, medium, and high values of cortical thickness correspond to the first quartile (25 percentile; red line), median (green line), and third quartile (75 percentile; blue line). The x-axis shows the timepoints where 0 corresponds to baseline, 1 to the year 1 follow-up, and 2 to the year 2 follow-up. The y-axis shows the estimated marginal means of standardized externalizing symptom scores. Panel A shows the model where baseline cortical thickness of the left pars triangularis predicted the trajectory of externalizing symptoms over the two-year follow-up period. Panel B shows the model where baseline cortical thickness of the left rostral middle frontal gyrus predicted the trajectory of externalizing symptoms over the two-year follow-up period.

## Discussion

In the present study, we examined cross-sectional and longitudinal relationships between grey matter structure in fronto-limbic/striatal brain regions and externalizing symptoms, CU traits, and emotion dysregulation in a large population-based sample of children. The results found were not completely consistent with our hypotheses. Although the three externalizing psychopathology dimensions all showed significant relations with fronto-limbic/striatal regions, either at baseline or over follow-up, specific brain alterations were found to be significant within each dimension. Emotion dysregulation scores were significantly and negatively associated with subcortical volume of limbic and striatal regions (greater impairment associated with lower volume). CU traits were significantly positively associated with frontal cortical thickness (higher traits associated with increased thickness). Greater baseline frontal cortical thickness was associated with greater decreases in externalizing symptoms over the two-year follow-up period. Overall, the results of the current work highlight the importance of examining the neurobiological correlates of different externalizing psychopathology dimensions given that they may be predictors of unique impairment pathways across development [51,52].

Greater emotion dysregulation was negatively associated with subcortical volume in the left caudate, right amygdala, and left and right nucleus accumbens, consistent with prior work [53,54]. These findings of the current study converge with prior work. Normative maturation of limbic and striatal structures has been linked to greater emotion regulation and related executive functions (such as inhibitory control [55]). Conversely, lower volume of striatal structures has been linked to increased irritability, both cross-sectionally [54] and longitudinally [53]. Notably, decreased or increased volume of various brain regions has been linked to hypo- or hyperactivity of those regions, respectively [56,57]. Although speculative, a potential explanation of the link between lower subcortical volume and emotion dysregulation is a possible hypoactivity of those regions impairing normative maturation of top-down modulation of prefrontal cortical regions; a pathway implicated in adaptive emotion regulation [58–60].

Elevated CU traits at baseline were positively associated with cortical thickness in the right rostral middle frontal gyrus and the left and right pars orbitalis. These brain regions have been associated with higher order cognitive functions, regulatory skills, and cognitive control [61]. Although it may seem counterintuitive that greater CU trait impairment is associated with increased thickness in these key regions, previous findings on this association have been mixed in terms of presence and direction of effects (for a review see [62]). One potential reason for the inconsistent findings is the different age ranges examined across prior studies. Many prior studies have focused on adolescent and young adult samples, as this developmental period generally coincides with the onset of more severe conduct problems [63,64]. The narrow and younger age range of the ABCD baseline sample (9-11 years old) overlaps with the developmental timing of cortical thickness transitioning from reaching peak thickness to reduced thickness due to mechanisms such as synaptic pruning [65–67]. This transition period may serve as a developmental window whereby differing patterns of structural development could potentially increase or decrease the risk of externalizing psychopathology. It is possible that at the cross-sectional level of analysis (during pre-adolescence), the increased cortical thickness linked to CU traits reflects delayed maturation of these regions (e.g., delayed synaptic pruning; [68].

We found longitudinal relationships between externalizing symptoms and cortical thickness such that participants in the highest quartile of baseline thickness of the pars triangularis and rostral middle frontal gyrus showed the greatest decreases in externalizing symptoms at the 1 and 2-year follow up timepoints, showing some consistency with a prior study [51]. The results of the longitudinal analysis differ in direction than the cross-sectional analysis between CU traits and cortical thickness. Both analyses implicate the rostral middle frontal gyrus being linked to these externalizing psychopathology dimensions. Although speculative, it is possible that higher cortical thickness at baseline reflects delayed maturation such that it is linked to higher externalizing psychopathology dimension impairment (CU traits), but it is followed by rapid maturation throughout adolescence leading to reduced cortical thickness and as a result, lower externalizing symptoms [69].

Variation in symptom endorsement between the three externalizing psychopathology dimensions across time may be contributing to the pattern of brain-behaviour relationships identified in the current study. Although each dimension showed different patterns of significance in cross-sectional brain-behaviour relationships, there were similar directional trends across all three (*Figure 1*) suggesting that frontolimbic and striatal regions are implicated across externalizing psychopathology dimensions. This is consistent with recent work suggesting that while neuroanatomical correlates of psychopathology may be dimension-, diagnostic-, or individual-specific, there may be overlapping functional networks implicated across similar dimensions or diagnoses [70]. Notably, it is possible that specific windows of child development are more likely to be characterized by unique dimensions of externalizing psychopathology compared to others. This would lead to greater variation of symptom scores required to detect a significant association with neurobiological metrics. For example, emotion dysregulation had the largest endorsement in the baseline sample with 11.9% of the sample showing no symptom endorsement compared to 59.3% for CU traits and 80.7% for externalizing symptoms. The greater endorsement of emotion dysregulation at baseline may have provided enough variation in symptoms to detect a significant cross-sectional effect with subcortical volumes. Further, the significant longitudinal relationship may be due to the overall decrease in externalizing symptom scores in the 1-year and 2-year follow up compared to the baseline sample; a pattern not found with emotion dysregulation or CU trait scores (see *Table 1*). This may have increased the variation of scores between timepoints, and thus, the likelihood to detect a significant effect. The results of this study highlight the importance of using clinical measurements and psychopathology dimensions that account for developmental context, and developmentally specific symptom endorsement. Exploring the neurobiological links of a single externalizing dimension (e.g., one that captures the covariance of the three dimensions analyzed separately in this study), particularly in a large public dataset, may not capture the full range of variation of externalizing psychopathology that can inform meaningful brain-behaviour relationships.

## Limitations

This study has several important strengths, but some limitations must also be considered. First, the measurement instruments used to assess symptoms and dimensions of psychopathology were limited to those included in the ABCD protocol. For example, the tool used to measure CU traits was composed of selected items taken from the CBCL and the SDQ. Although this tool has previously been used to investigate CU traits in the ABCD sample [44], a measure specific to CU traits such as the Inventory of Callous Unemotional Traits [71] might have indexed more specific and validated CU traits. Second, the sample is limited by a narrowly defined age range. This narrow age range allows for the investigation of a specific developmental window implicated in brain maturation [29] pre-clinically significant externalizing psychopathology [62]. However, it is important to expand beyond this age range to strengthen our understanding of neurobiological differences for various externalizing psychopathology dimensions across the lifespan. Third, although we did not specifically examine race in our analyses, the year-2 sample features a substantial increase of no-response participants when asked about race compared to baseline and year-1 (see *Table 1*). Thus, the generalizability of the current results to the overall pediatric population in the U.S. may be limited. Fourth, we focused the scope of the current study to understanding whether different dimensions of externalizing psychopathology have different brain structure correlates. However, we acknowledge the possibility of these relationships being influenced by environmental outcomes or early life events [72–74]. Fifth, externalizing symptoms in the current study represented a total count of symptoms from ODD and CD traits from the K-SADS which includes severe conduct behaviors that may typically begin to emerge in late adolescence (e.g., breaking and entering, vandalism; [63]) leading to low endorsement in the ABCD baseline sample. Finally, while the large mainly normative sample allowed us to examine the full range of externalizing psychopathology symptom range present in the general population [46], it is possible that more significant cross-sectional and longitudinal brain-behaviour relationships would be found in a sample enriched for externalizing psychopathology.

## Conclusion

Taken together, our findings indicate that while brain structures implicated in the fronto-limbic/striatal networks are linked to externalizing psychopathology severity across a pediatric population-based sample, each dimension and the time-point being measured may influence the pattern of brain-behavior relations found. Longitudinal findings suggest that regional brain structure in middle childhood may influence externalizing symptoms measured two years later. Future research incorporating additional outcome variables, environmental metrics, such as parent and caregiver relationships [75], and in clinically enriched samples might shed further light on brain signatures that influence better versus worse outcomes among children over development.

## Acknowledgements

Data used in the preparation of this article were obtained from the Adolescent Brain Cognitive Development® (ABCD) Study (https://abcdstudy.org), held in the NIMH Data Archive (NDA). This is a multisite, longitudinal study designed to recruit more than 10,000 children age 9–10 and follow them over 10 years into early adulthood. The ABCD Study® is supported by the National Institutes of Health and additional federal partners under award numbers U01DA041048, U01DA050989, U01DA051016, U01DA041022, U01DA051018, U01DA051037, U01DA050987, U01DA041174, U01DA041106, U01DA041117, U01DA041028, U01DA041134, U01DA050988, U01DA051039, U01DA041156, U01DA041025, U01DA041120, U01DA051038, U01DA041148, U01DA041093, U01DA041089, U24DA041123, U24DA041147. A full list of supporters is available at https://abcdstudy.org/federal-partners.html. A listing of participating sites and a complete listing of the study investigators can be found at https://abcdstudy.org/consortiummembers/. ABCD consortium investigators designed and implemented the study and/or provided data but did not necessarily participate in analysis or writing of this report. This manuscript reflects the views of the authors and may not reflect the opinions or views of the NIH or ABCD consortium investigators. The ABCD data repository grows and changes over time. The ABCD data used in this report came from NDA Release 4.0 (DOI: 10.15154/q36k-ga33).

## Author Contributions

LP and HN conducted all the analyses and wrote the main draft of the paper. BA, ACB, and SA contributed to the design of the study. MS contributed to the design of the analytical approaches used. All authors wrote, revised, approved, and agreed to be accountable for all aspects of the final manuscript.

