## Supplementary Materials for "Investigating cross-sectional and longitudinal relationships between brain structure and distinct dimensions of externalizing psychopathology in the ABCD Sample"

Figure S1. Study Flow

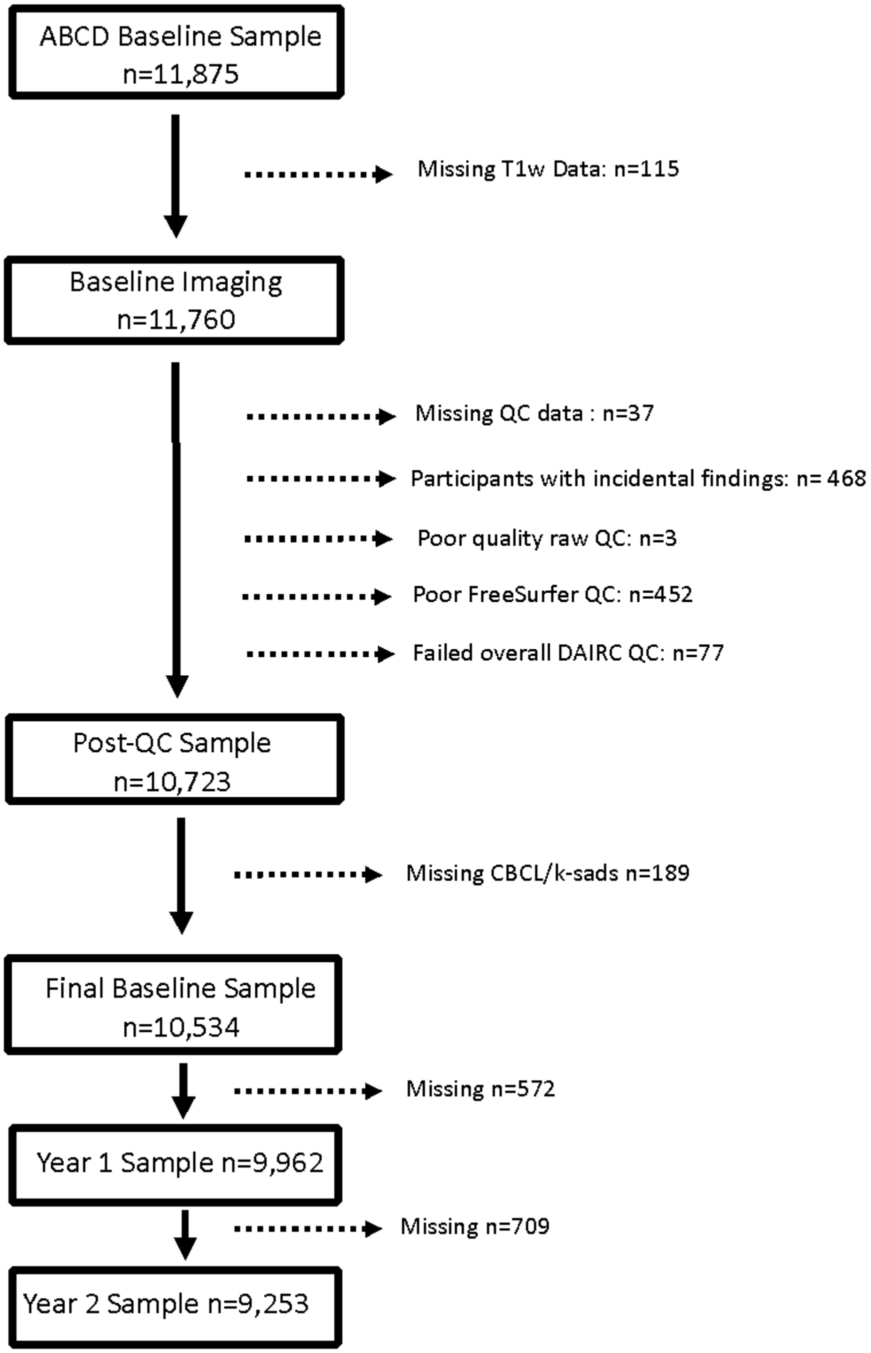


Note: The variables used to exclude participants are as follows. The incidental findings variable was mrif_score and participants were excluded if they were considered to "need clinical referral" or "immediate clinical referral". Participants with a zero value for the iqc_t1_ok_ser variable were excluded indicating that they had poor quality raw T1-weighted scans. Participants were excluded if they received a "reject" from the fsqc_qc variable indicating they failed FreeSurfer QC. Finally, any additional participants who failed the DAIRC QC were excluded (received a zero for the imgincl_t1w_include variable).

Table S1. ROIs used in Present Study Based on Prior Literature

| Region | Emotion Dysregulation | CU Traits |
|---|---|---|
| **Subcortical** | | |
| Amygdala | (Seymour et al., 2017) | (Cardinale et al., 2019; Gao et al., 2020; Ibrahim et al., 2021; Waller et al., 2020) |
| Caudate | | (Fairchild et al., 2013) |
| Putamen | (Seymour et al., 2017) | |
| Nucleus Accumbens | | (Cha et al., 2015) |
| **Cortical** | | |
| Insula | | (Chaarani et al., 2020; Sterzer et al., 2007; Waller et al., 2020) |
| Anterior Cingulate Cortex (rostral and caudal) | (Mulraney et al., 2021) | (De Brito et al., 2009; Sterzer et al., 2007) |
| Inferior Frontal Gyrus (pars opercularis, pars triangularis, pars orbitalis) | (Chaarani et al., 2020) | (Cha et al., 2015) |
| Middle frontal cortex (rostral and caudal) | (Mulraney et al., 2021; Tsai et al., 2020) | |

Table S2. Full ROI Labels

| ABCD ROI Label | Full ROI Label |
|---|---|
| Cortical Thickness | |
| R_RostMidFrontal | rostral middle frontal (right hemisphere) |
| L_RostMidFrontal | rostral middle frontal (left hemisphere) |
| R_RostAntCingulate | rostral anterior cingulate (right hemisphere) |
| L_RostAntCingulate | rostral anterior cingulate (left hemisphere) |
| R_ParsTriangularis | pars triangularis (right hemisphere) |
| L_ParsTriangularis | pars triangularis (left hemisphere) |
| R_ParsOpercularis | pars opercularis (right hemisphere) |
| L_ParsOpercularis | pars opercularis (left hemisphere) |
| R_ParsOrbitalis | pars orbitalis (right hemisphere) |
| L_ParsOrbitalis | pars orbitalis (left hemisphere) |
| R_insula | insula (right hemisphere) |
| L_insula | insula (left hemisphere) |
| R_CaudMidFrontal | caudal middle frontal (right hemisphere) |
| L_CaudMidFrontal | caudal middle frontal (left hemisphere) |
| R_CaudAntCingulate | caudal anterior cingulate (right hemisphere) |
| L_CaudAntCingulate | caudal anterior cingulate (left hemisphere) |
| Subcortical Volume | |
| R_Putamen | putamen (right hemisphere) |
| L_putamen | putamen (left hemisphere) |
| R_caudate | caudate (right hemisphere) |
| L_caudate | caudate (left hemisphere) |
| R_amygdala | amygdala (right hemisphere) |
| L_amygdala | amygdala (left hemisphere) |
| R_accumbens | accumbens area (right hemisphere) |

| | |
|---|---|
| L_accumbens | accumbens area (left hemisphere) |

Note. Cortical thickness in mm of APARC ROI; Volume in mm^3 of ASEG ROI
